# Partial sleep deprivation diminishes and uncouples neuronal activity and energy metabolism in the live mouse hippocampus

**DOI:** 10.64898/2026.09.25.754485

**Authors:** Zhengyi Lu, Kevin Mai, Alfredo Cardenas-Rivera, Chang Liu, Yuntao Li, Priya Shah, Kai Zhang, Bryan Q. Spring, Mohammad A Yaseen

**Author notes:** **Corresponding Author:** Mohammad Abbas Yaseen, Department of Bioengineering, Northeastern University, 360 Huntington Ave, ISEC 206, Boston, MA USA 02115.

## Abstract

Sleep disorders and wakefulness constitute a pronounced and pervasive health risk that contributes to numerous cerebral pathologies such as stroke, depression, and neurodegenerative disorders. The hippocampus (HC) plays vital roles in learning and memory and is especially susceptible to the pathological insults imposed by sleep deprivation (SD). While SD disrupts neuronal network signaling and triggers inflammatory processes, the impact of SD on metabolism within the HC presently remains poorly understood. In this study, we assessed the progressive and cumulative effects of partial SD in the live mouse HC CA1 region using two-photon microscopy. Using a custom-designed implantable cannula enabling direct HC visualization and administration of dyes and pharmacological agents directly to HC tissue, we longitudinally monitored spontaneous neuronal activity and partial pressure of oxygen (pO₂) within HC CA1 tissue before and during a 7-day period of partial SD and after a 7-day period of recovery. Our results revealed a progressive decline in spontaneous neural activity, alongside sustained elevation in tissue pO₂ during the 7-day partial SD period. Notably, after 7 days of recovery, neuronal activity did not return to baseline, and tissue oxygenation continued to increase. Our findings emphasize chronic SD’s cumulative and enduring impairments to neuronal function and metabolism in the HC and reveal uncoupling between metabolic and functional recovery that outlasts the sleep restriction period.

## Introduction

Sleep disorders and wakefulness impact hundreds of millions of people globally and notably augment risks for cardiovascular disease, obesity, diabetes, hypertension, and mental health challenges. [1–3]. In a systematic review and meta-analysis including 1.1 million individuals, insufficient sleep was reported in approximately 25% of the populations within the Netherlands, United Kingdom, and the United States [4]. Worldwide, another review estimated global incidence rates of 16.2% and 7.9% for insomnia and severe insomnia in adults [5]. In the brain, sleep disturbance is implicated in cognitive and memory deficits and the advancement of stroke, depression, and dementia-related neurodegenerative disorders [6,7]. The hippocampus (HC) makes indispensable contributions to memory formation and consolidation, cognition, and navigation, and is especially vulnerable to disturbances like sleep deprivation (SD) and hypoxic insult [8,9]. During sleep, memory consolidation is strengthened by conditioning excitatory and inhibitory synaptic connections and through regulation of neuronal firing rates between and within the HC and cortex [10–12] In animal models, SD has been shown to induce reductions in HC synaptic proteins like postsynaptic protein 95 (PSD-95) and synaptophysin (Syn), and phosphorylation activity of memory-related kinases such as extracellular signal-regulated kinase (ERK) [13,14]. In human subjects, SD has been shown to alter functional connectivity within the HC, and altered functional connectivity between the HC and frontal gyrus has been strongly correlated with severe insomnia and diminished sleep quality [15,16]. SD has also reportedly facilitates neurodegeneration by aggravating multiple pathological processes, including increased oxidative stress, neuroinflammation, and impaired protein clearance [6].

To date, the impacts of SD on energy metabolism within the HC remain poorly understood [17]. These current gaps in our knowledge limit clinical characterizations of SD’s disruptions using translational neuroimaging technologies like fMRI and PET. Moreover, mechanistic interpretations are currently lacking to explain the limited capacity of recovery sleep to counteract the effects of SD [18–21]. To gain more detailed insight into the adverse pathological effects of SD, and why these effects are especially persistent within the HC, advanced tools are required that enable (1) longitudinal assessment of the live, intact HC with cellular resolution and (2) direct administration of contrast agents and pharmacological treatments to the HC tissue.

In this study, we longitudinally evaluated the impact of SD on spontaneous neuronal signaling and the partial pressure of oxygen (pO₂) in the live mouse HC CA1 region using intravital two-photon microscopy (2PM). After designing an implantable cannula to enable both gradient-index (GRIN)-lens mediated optical microscopy and direct drug administration to subcortical brain structures, we visualized spontaneous activity in CA1 neurons and tissue pO₂ in transgenic Thy1-GCamp6f mice over the course of a 7-day period of partial SD and after 7 days of recovery. We specifically employed a partial SD paradigm to mimic the chronic insufficient sleep resulting from habitual late bedtimes or other lifestyle-induced sleep insufficiency commonly observed in modern society [22]. Over the course of the 7-day partial SD period, we observed reductions in spontaneous calcium events in CA1 neurons and concurrent increases in tissue pO₂. After a 7-day recovery period of normal sleep and diet, neuronal activity did not return to baseline levels and tissue pO₂ increased slightly higher. To our knowledge, we demonstrate the first high-resolution oxygen mapping within subcortical brain structures of the live brain. Our study highlights the cumulative burden imposed by chronic partial SD on hippocampal neuronal signaling and tissue metabolism and a dissociation between metabolic and functional recovery that outlasts the sleep restriction period. The results will help improve our understanding of the HC’s heightened vulnerability to sleep insufficiency and SD’s role in the onset and progression of neurodegenerative disorders.

## Methods

### Cannula design and validation

To complete this intravital, longitudinal assessment in subcortical brain tissue, we first designed and validated a custom GRIN lens cannula that enabled both high resolution imaging and direct administration of exogenous contrast and pharmacological agents. Our cannula extends the utility of a prior design by Bocarsly et al that prioritized biocompatibility, minimal displacement of brain tissue, and longitudinal imaging capabilities [23]. Like Bocarsly et al’s original design, the wall of our cylindrical cannula was made from biocompatible polyimide tubing with 1.17 mm outer diameter and 25.4 μm shell thickness (Microlumen, model: 440-I) to minimize displacement of brain tissue (Figure 1.a). We altered the window material and the fabrication technique to enable repeatable tissue injections with minimal gas exchange. The injection-permeable window material, Flexdym (Eden Tech), is a biocompatible and optically transparent thermoplastic elastomer. Compared to more common materials like Polydimethylsiloxane (PDMS), Flexdym exhibits lower gas permeability and lower adsorption of small molecules, which helps maintain integrity of the brain tissue environment. Crucially, the Flexdym material’s ‘self-healing’ capacity enables post-injection imaging [24,25]. Details for the cannula construction and validation are provided in the supplementary materials.

**Figure 1.**
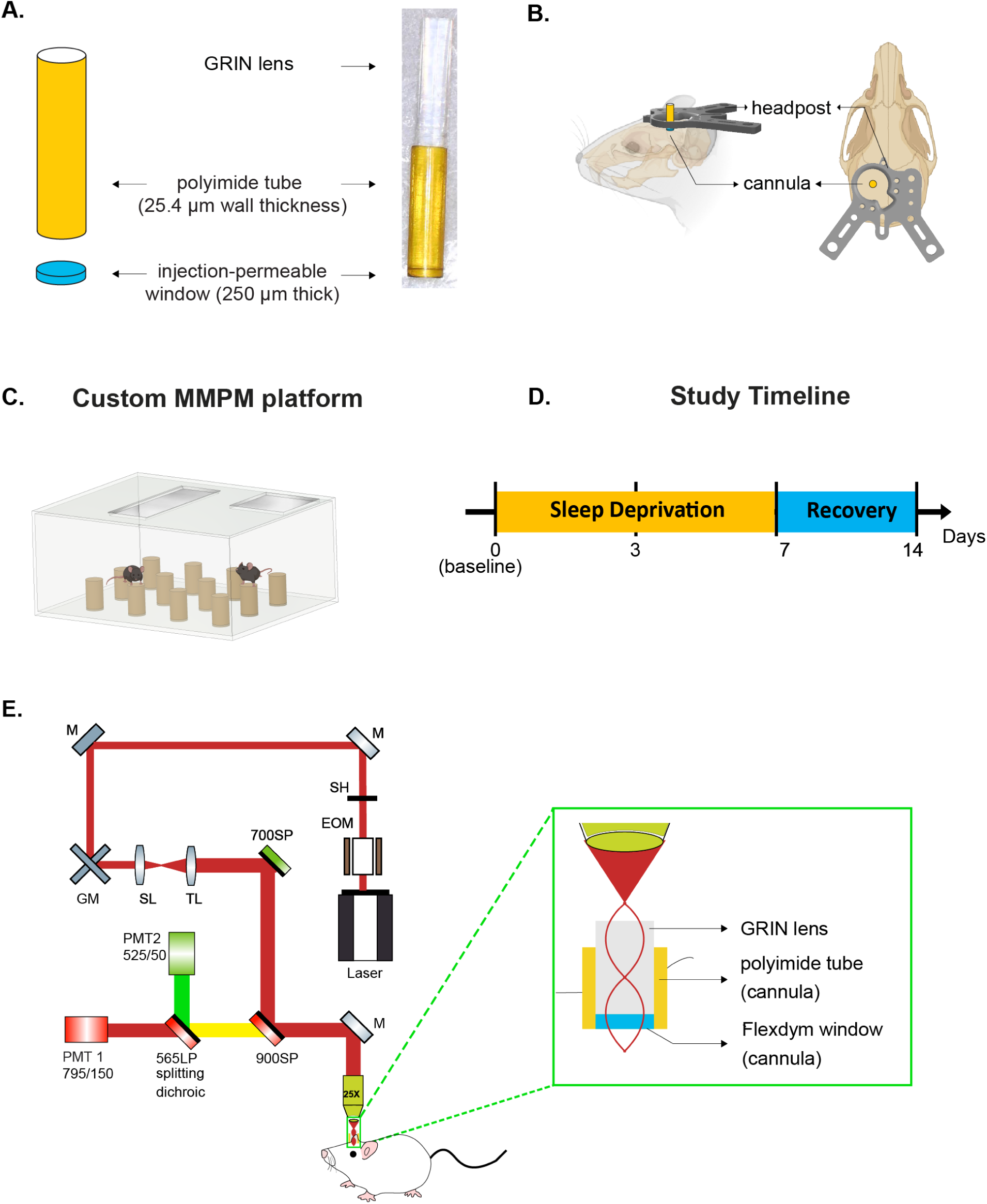
(a) Schematic of the custom cannula (left) and photograph of the assembled cannula with the GRIN lens (right). (b) Implantation of the cannula into the mouse HC and custom headpost installation. (c) Custom MMPM platform. Platform diameter: 3 cm; height: 4.5 cm; spacing: 5cm. The water level was maintained 1 cm below the platform surface. 3-4 mice were usually housed in the MMPM chamber simultaneously. (d) Study timeline. Measurements were performed on Days 0, 3, 7, and 14. (c) Two-photon imaging system diagram. Imaging was performed under 950 nm excitation. PMT1 was used for tissue oxygen measurements using Oxyphor 2P (ex: 950 nm; em: 735–775 nm). PMT2 was used for neuronal calcium imaging using GCaMP6f (ex: 920–950 nm; em: 510–515 nm). Mouse illustrations in (b) and (c) were created using assets from BioRender.com.

### Animal preparation

All procedures involving animals adhered to ARRIVE guidelines and were approved by the Animal Care and Use Committee at Northeastern University. Eight female C57BL/6J-Tg (Thy1-GCaMP6f) GP5.17Dkim/J mice (age 9-11 months, Jackson Laboratory, # 025393) and twelve age-matched wildtype cagemates were used in this study. In each SD cycle, 1-2 Thy1-GCaMP6f mice were paired with 1-3 WT cagemates. These transgenic mice express the fluorescent calcium indicator, GCaMP6f, primarily in subsets of excitatory neurons in the brain under Thy1 promoter. Fluorophore expression is stable over months across multiple brain regions, enabling long-term imaging with high signal-to-noise ratio in behaving mice [26]. Mice underwent cannula implantation surgery and were allowed to recover for ∼ 6 weeks.

Cannula implantation followed a surgical protocol established previously for GRIN lenses (Figure 1.b) [27]. Under isoflurane anesthesia, the skull was exposed by hair and skin removal. Using bregma as the reference point, the skull was marked with a 1mm diameter circle (AP:-2mm ML:1.8mm to bregma), and then drilled using a 0.3-mm drill bit. The drilled bone fragment was removed with fine forceps. The dura was pierced by a 31G needle, and the cortical tissue above HC was gently aspirated using a 25G blunt needle and custom-built suction system (Figure S1.a). Aspiration continued until the corpus callosum became visible. The cannula was thoroughly sterilized with 70% ethanol and inserted into the brain using a custom-built, vacuum-mediated holder. Using a portable digital microscope for guidance, the cannula was lowered into the brain to an approximate depth of ∼1650μm from the skull surface at 5 μm/s (Figure S1.b). Once the cannula was positioned, it was secured to the skull with Loctite 4014 adhesive. A custom headpost was then secured to the skull using adhesive (Loctite 401). To minimize cerebral inflammation and edema, Dexamethasone (4 mg/ml, 4.8 mg/kg) was administered subcutaneously approximately 4-8 hours before surgery. Saline-soaked gel foam was used to control any bleeding during the procedure. A heating pad was used to maintain the animal’s body temperature. Post-operative medications were administered in accordance with institutional guidelines. At times when the animal was not imaged, a custom cap was fixed to the headpost to cover the cannula, protecting the cannula from dust or mechanical damage.

Habituation training for head fixation initiated after a 2-3 week surgical recovery. During training, mice were secured to the cradle by fastening the headpost to the cradle frame. For positive reinforcement, mice were given small amounts of sweetened condensed milk at intervals. Initial training sessions lasted for 10 minutes, with the duration increased by 10 minutes each day until mice could remain calmly head-fixed for up to one hour.

### SD protocol

Figure 1.c-e displays the salient features of our longitudinal SD study. After surgery and acclimation training, mice were subjected to 7 days of partial SD. Tissue pO_2_ and Thy1 neuronal activity were measured in CA1 via 2PM on Days 0 (baseline), 3, 7, and 14 (following 7 days of recovery). A within-subject design was employed, in which the baseline measurement of each mouse served as its own control. When not subjected to partial SD, the mice were housed under 14h:10h light/dark cycle (light on at 7AM, off at 9PM) and received food and water ad libitum.

The modified multiple platform method (MMPM) was utilized to implement partial SD (Figure 1.c). MMPM is a widely-utilized model that completely abolishes rapid eye movement (REM) sleep and decreases non-REM (NREM) sleep by approximately 35-40%. It was designed to prevent movement restriction and attenuate stress from social isolation [28–31]. In our design, 12 small platforms (diameter: 3 cm, height: 4.5 cm, spacing: 5cm) were arranged in a water tank containing 3-4 mice. The tank was filled with water maintained at 27 °C using a heating pad positioned below the chamber. The water level was maintained 1 cm below the platform surface, and the water was changed daily for sanitary purposes. The mice remained in the MMPM chamber for 18 hours daily (from 4 PM to 10 AM of the next day) with continued access to food and water ad libitum. All mice were returned to their home cages from 10 AM to 4 PM. The partial SD protocol was administered for 7 days (Figure 1.d). Wildtype cagemates were included in the MMPM chamber to provide social companionship. To prevent excessive weight loss and to maintain adequate hydration and nutritional status during the SD period, the mouse diet was supplemented with DietGel® Recovery (ClearH2O), a nutritionally fortified water-based gel. Following completion of SD, mice were allowed a 7-day recovery period under standard housing conditions, with normal light–dark cycles and ad libitum access to food and water. Body weight was measured daily at 4 PM for all mice, starting on the first day of SD and continuing for 40 days.

### In vivo imaging

All measurements were performed using a commercial two-photon microscope (Bruker, model: Ultima2P plus), a tunable ultrafast pulsed laser with a repetition rate of 80 MHz (Spectra-Physics, model: InSight DeepSee X3+), a water-immersion 25X objective (Nikon, model: TH-N25X-APO-MP), and a GRIN lens (Thorlabs, model: G1P11). Before imaging, the GRIN lens was pre-warmed to approximate body temperature. To maintain physiological temperature and reduce temperature-related variability, the objective was heated using an objective heater. A feedback-based temperature controller (ALA Scientific Instruments) was used to prevent overheating of the objective. During imaging, the mouse was secured onto the cradle, and the GRIN lens was gently inserted into the cannula. Water was added between the objective and the GRIN lens for optical coupling. Neuronal GCaMP and hippocampal tissue pO_2_ measurements were performed using an excitation wavelength *λ* = 950 nm. During the imaging experiments, mice were periodically rewarded with sweetened condensed milk between measurements.

#### Neuronal imaging

To assess spontaneous CA1 neural activity, time-series recordings of GCaMP fluorescence were acquired within 151.4 × 151.4 µm field of view (FOV) at 10.1 Hz for 8 min. At the initial imaging session, the FOV was chosen to include the largest number of clearly identifiable neurons. Using microvascular morphology as fiducial markers, the same FOVs were chosen, to the best of our ability, across time points. If the number of active or identifiable neurons within the original FOV became insufficient for analysis over the course of SD, an additional FOV containing a high neuronal density was acquired in the same session. For subsequent imaging days, both the original FOV and any supplementary FOVs were re-acquired whenever possible. During data analysis, priority was always given to the FOV that matched the baseline session; supplementary FOVs were analyzed only when the baseline-matched FOV contained too few identifiable neurons for reliable analysis.

#### Tissue oxygen imagin

PO_2_ was mapped with high resolution in hippocampal tissue using the phosphorescent oxygen sensitizer, Oxyphor 2P. Oxyphor 2P is a phosphorescent metalloporphyrin dye meticulously designed for absolute pO_2_ quantitation via 2PM [32]. Briefly, oxygen molecules quench the phosphorescent emission originating from the excited triplet state Oxyphor 2P’s metalloporphyrin core. The molecule’s phosphorescence lifetime follows the Stern-Vollmer relationship with ambient oxygen in the surrounding microenvironment. A quantitative readout of the local pO_2_ is provided by a calibration curve [32].

After neuronal activity measurements, the GRIN lens was removed from the cannula for Oxyphor 2P injection. Using a motorized micromanipulator and syringe pump, ∼ 120 nL of Oxyphor 2P (340 μM solution) was delivered at 10 nL/s. The 34G Hamilton injection needle was advanced at 5 μm/s, lowered approximately 1500 μm beyond the bottom of the cannula, and then retracted by 900 μm.

Following injection, the Oxyphor 2P was allowed to diffuse into CA1 tissue for at least 1 h to ensure sufficient distribution throughout the imaging field. Under 950 nm excitation, the imaging plane was selected as one that displayed distinct vascular morphology vascular pattern and the largest available region of Oxyphor 2P signal. PO₂ measurements were acquired in a grid pattern. Each grid typically consisted of points spaced 30-40 μm apart, yielding approximately 230-340 measurement points per mouse. To minimize potential thermal effects from repeated excitation, the measurement locations were scanned in random order. At each point, two-photon phosphorescence lifetime imaging microscopy was repeated three times, with each repetition consisting of 1000 excitation–detection cycles. In each cycle, a 10 μs excitation pulse was followed by a 290 μs photon collection window.

### Body weight assessment

As a simple assessment of partial SD’s systemic impact, body weight was measured daily from day 0 to day 40 in 3 cohorts of animals: (cohort 1) the animals that underwent surgical cannula implantation and 7 days partial SD and imaging, (cohort 2) animals that underwent only 7 days partial SD, and (cohort 3) control animals without surgical implantation and without SD. Animals cohorts that underwent partial SD maintained ad libitum access to water and food, including a DietGel® Recovery supplement, during the 7-day partial SD period. After the 7-day partial SD period, all animals returned to normal diet and light/dark cycle.

#### Data Analysis, Neuronal Activity

Transient recordings of neuronal GCaMP fluorescence were processed in MATLAB using a custom-built pipeline based on CaImAn, an open-source toolbox designed for large-scale single- and two-photon calcium imaging analysis. CalmAn provides automatic and scalable methods for motion correction, neuronal source extraction, and calcium signal denoising, generating the calcium event trace for each detected neuron [33].

##### Single neuron activity

To assess spontaneous neuronal activity at the single neuron level, we analyzed four features of the calcium transients: firing count, mean event amplitude (ΔF/F), mean event duration, and mean event area under the curve (AUC), computed by averaging across all events detected in that neuron. The CalmAn software segments and ranks neurons based on a combination of their spatial footprints and their temporal activity [34]. To avoid selection bias and standardize analysis across all measurements, single neuron activity was assessed by analyzing the ten most highly ranked neurons in each measurement. For each neuronal calcium trace, a baseline intensity segment was manually selected, after which preliminary peak detection was performed using two thresholds: the minimum peak height, defined as baseline mean plus two baseline standard deviations; and the minimum peak prominence, defined as eight times the baseline standard deviation. In subsequent large-scale data processing, random spot-checks were performed to confirm that peak detection remained reliable with the established thresholds. To identify calcium events, we defined the threshold as 5% change from the baseline [35]. An event was identified as any continuous period during which the calcium signal rose above this threshold and subsequently returned below it. Firing count was computed as the total number of detected events within the recording period. For each event, only peaks that passed the previously defined peak-detection criteria were considered, and only the highest peak within each event was retained for analysis. Event amplitude was defined as the ΔF/F value of the retained peak. Event duration was defined as the interval between the signal crossing above and falling back below the event threshold. Area under the curve (AUC) was calculated by integrating the ΔF/F trace over the event duration.

##### Network activity

Coordinated neuronal activity was evaluated at the population level by quantifying synchronous, spontaneous firing. Calcium event onset frames were detected by CaImAn and converted into binary event trains, in which event-onset frames were assigned as 1 and all other frames as 0. To accommodate temporal jitter in event detection, each onset was expanded into a seven-frame window (±3 frames). The percentage of co-activated neurons at each sampling frame, denoted as *φ*(*t*), was calculated as the number of simultaneously firing neurons divided by the total number of neurons in the original dataset [36]. Spectral composition of *φ*(*t*) was evaluated by its power spectral density (PSD). The spectral power beyond 1 Hz band was minimal with little meaningful signal and was therefore excluded, with subsequent analysis focused on the 0-1 Hz band. [36–38]. The resulting power spectrum was smoothed using a five-point moving-average window.

#### Data Analysis, Tissue oxygenation

Using custom-written MATLAB tools, hippocampal tissue pO_2_ values were quantified by calculating the lifetimes of time-resolved Oxyphor2P phosphorescence decays. The first 5 μs of each phosphorescence decay trace was discarded to minimize the influence of the system response function, leaving 285 μs of usable decay per cycle [32]. For each repetition, photon counts from all 1000 cycles were accumulated to produce a decay curve. The three resulting decay curves were visually inspected to identify potential outliers, and if no anomalies were found, the curves were averaged to generate the final decay trace for that location, from which the pO₂ value was calculated. The final decay was fitted to an empirical bi-exponential function, *pO*_2_ = *a*_0_ + *a*_1_*e*^−τ/*a*3^ + *a*_2_*e*^−τ/*a*4^, where τ is the phosphorescence lifetime, a_0_-a_4_ are empirically determined fitting coefficients. To improve fitting accuracy and ensure adequate photon statistics, any measurement point with fewer than 7500 accumulated photons was excluded from analysis.

### Statistical analysis

All statistical analyses were performed using both custom-written and publicly-available MATLAB scripts, as described below.

For neuronal activity at the individual level (firing count, ΔF/F, event duration, AUC): hierarchical bootstrap (HB) was applied to estimate confidence intervals for individual timepoints and to obtain p-values for pairwise timepoint comparisons. HB is particularly well-suited for handling hierarchical data structures, most commonly the activity of multiple neurons in multiple animals across multiple trials, and has been increasingly applied in high-level neuroscience studies [20,39]. For each analysis, 10,000 bootstrap resamples were performed.

To evaluate number of neurons per FOV and tissue pO₂, Friedman test was first applied to assess whether the measured variables differed across timepoints. If the Friedman test indicated a significant overall effect, pairwise comparisons were performed using the Wilcoxon signed-rank test. Both tests are designed for repeated-measures nonparametric data, suitable for small sample sizes [40]. To control the false discovery rate (FDR) across multiple comparisons, p-values were adjusted using the Benjamini-Hochberg (BH) procedure with an FDR threshold of 0.05.

Sample size was n = 7 or 8 for this exploratory study. For neuronal activity analysis, one mouse was excluded for exhibiting lower neuronal firing rates at baseline. To avoid undue influence from extreme outlier values, statistical analyses of tissue pO_2_ were performed by averaging the per-mouse medians at each timepoint [38]. Statistical significance was defined as p < 0.05 (*p < 0.05, **p < 0.01, ***p < 0.001). Analyses were primarily performed using custom-written code, with the hierarchical bootstrap component modified based on publicly available code [41].

## Results

Before, during, and after a prolonged period of partial SD, we imaged neuronal GCaMP activity and tissue pO_2_ in the live mouse CA1 region via 2PM through a GRIN lens. For 7 days, Thy1-GCamp6f mice remained on their normal light/dark cycle but were only able to sleep for a maximum of 6h each day (10:00AM – 4:00 PM). For the remaining 18h, the mice maintained their access to food and water ad libitum but were subjected to SD via a custom-built, heated multiple platform chamber. Based on prior studies, we employed this paradigm as a model for moderate, temporary partial SD and followed it by monitoring prospective recovery after 7 days. The approach was implemented to mimic a realistic and prevalent partial SD model for humans, particularly in urban environments [19,22].

### Cannula design and validation

Building on the developments of Bocarsly et al, we first designed an implantable, injection-permeable GRIN-lens cannula to enable both optical access and direct administration of contrast and pharmacological agents to the living rodent HC [23]. We evaluated the cannula’s fabrication process, structural viability, injection permeability, and imaging quality using commercial and custom imaging phantoms. We then optimized the dye injection protocol in live mice. Validation studies were performed in triplicate. All cannulae maintained their integrity and imaging viability after implantation and dye injection. Imaging quality through the cannulae was not affected by repeated needle penetration of the transparent endcap. Details are provided in the supplementary materials.

### Partial SD-suppresses neuronal activity, without recovery

Figure 2 displays the impact of partial SD on spontaneous activity within hippocampal Thy1 neurons. Consistent with prior reports, CA1 neurons demonstrated sparse, spontaneous activity with high population heterogeneity at day 0 (Figure 2.b) [42,43]. Over the 7-day SD period, the average number of spontaneous neuronal events progressively decreased, dropping nearly 50% from 25 to 13. The variability in neuronal firing rate, calculated by the firing rate’s coefficient of variation, also increased at day 7. From baseline (Day 0) to Day 3, CA1 neurons also demonstrated significant reductions in spontaneous event duration and AUC of spikes (Figure 2.b). The mean duration of neuronal events decreased from 2.1 s to 1.3 s from Day 0 to Day 3 and showed a slight increase afterward. However, the differences in duration did not reach statistical significance beyond Day 3. AUC followed a similar pattern, showing a significant decrease from baseline to Day 3, with modest, statistically insignificant increases at Day 7 and 14. The magnitude of spontaneous activity was unaffected by the partial SD insult. Normalized ΔF/F amplitudes remained consistent across all timepoints, with mean values remaining close to 1 and similar dispersity across timepoints (Figure S3.a). The average total number of active neurons per FOV dropped from approximately 24 at baseline to 20 at Day 3, 17 at Day 7, and recovered to about 21 at Day 14 (Figure S3.b).

**Figure 2.**
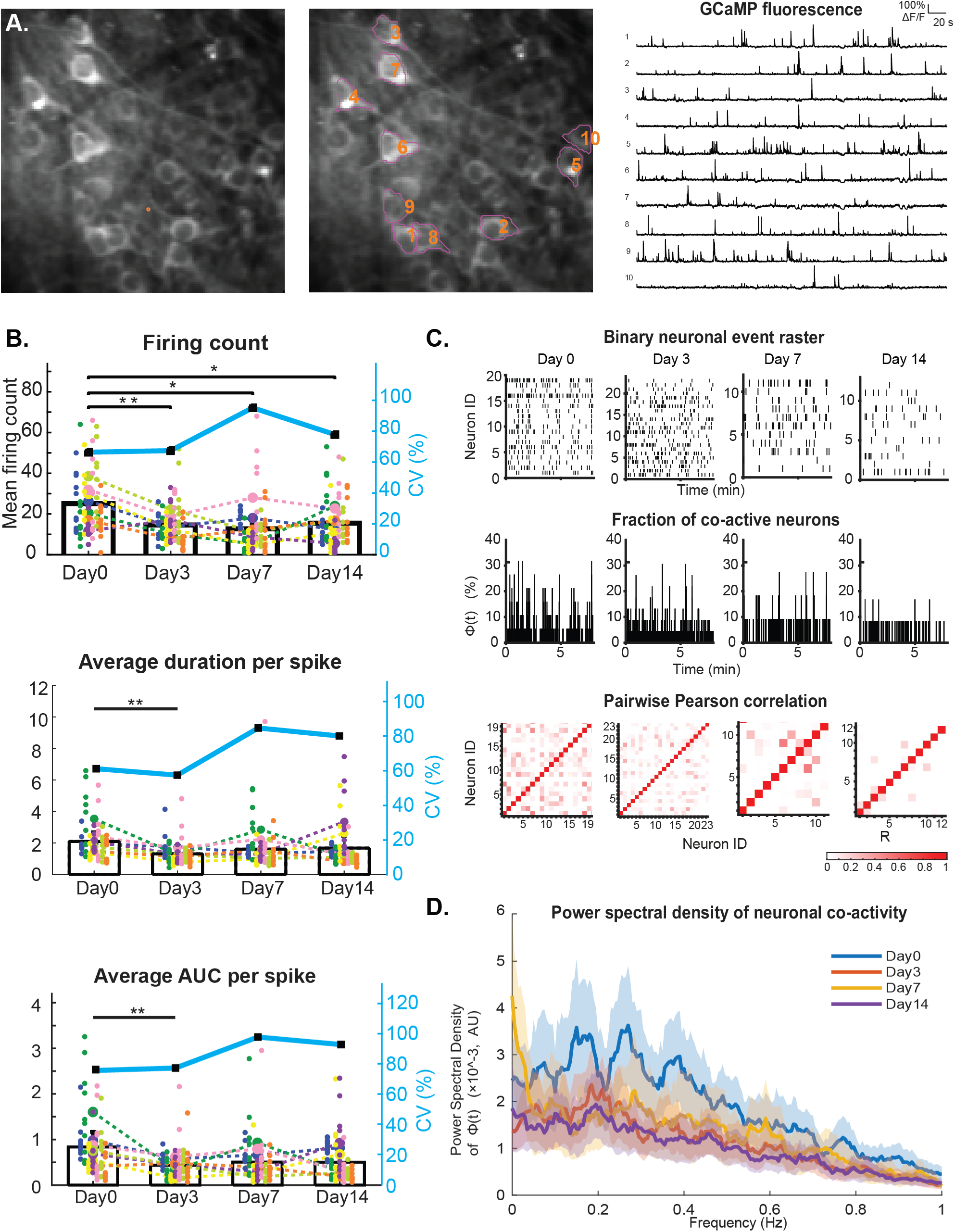
(a) Representative average intensity projection of CA1 neuronal imaging and corresponding calcium traces. Ten representative neurons identified by CaImAn and their corresponding calcium traces are shown. (b) Spontaneous neuronal firing count, spike duration, and area under curve, quantified before, during, and after partial SD. (c) Representative synchrony analysis from one animal. Top: binarized neuronal events rasters across the longitudinal SD study. Middle: fraction of co-active neurons (*φ*(*t*)) over time. Bottom: pairwise Pearson correlation matrices of neuronal activity. (d) PSD of *φ*(*t*) diminishes during partial SD and is not restored after 7 days recovery. (n=7)

To characterize coordinated network activity, the incidence of spontaneous calcium spikes were detected for all active neurons at all timepoints using CaImAn software and converted into binary event trains (Figure 2.c). The percentage of simultaneous neuronal events, *φ*(*t*), was calculated and evaluated following previously established methods [36–38].

During the 7-day SD event, we observed reductions in the spectral power of *φ*(*t*) (Figure 2.d). The PSD’s amplitude from 0 −1Hz diminished from Day 0 to Day 3 and remained low for the remainder of the SD period. While the power spectrum of *φ*(*t*) exhibited multiple pronounced peaks between 0.05 and 0.4 Hz at Day 0, the peaks were not evident after initiation of SD. Notably, we also observed reductions in the total number of spontaneously firing neurons (Figure S3.b), which lowers the denominator value for *φ*(*t*). Despite this reduction, total number of active neurons at each timepoint still provided sufficient statistical power for our analyses.

Strikingly, a 7-day recovery period of normal sleep and diet was insufficient to restore neuronal activity. At Day 14, both spontaneous firing count and correlated network activity remained suppressed after a recovery period that was equal in duration to the original SD insult.

### Partial SD induces elevations in hippocampal tissue pO₂, without recovery

By repeatedly administering the phosphorescent oxygen sensitizer, Oxyphor2P, directly to CA1 brain tissue, we assessed how hippocampal tissue pO_2_ varies in awake mice before, during, and after the 7-day partial SD insult [32]. To our knowledge, these are the first high-resolution observations of tissue pO_2_ from an intact, subcortical brain region obtained via two-photon phosphorescence lifetime microscopy. Prior to partial SD, hippocampal tissue pO_2_ displayed an average value of 20.6 ± 2.9 mmHg, lower than tissue pO_2_ observations around penetrating arterioles within the awake mouse somatosensory cortex [44]. During the partial SD period, we observed a significant, progressive increase in hippocampal tissue pO_2_ at every time point. Within individual mice, hippocampal pO_2_ values displayed skewed Gaussian distributions that progressively shifted higher at subsequent timepoints (Figure 3.a). Skewed Gaussian profiles were maintained after pooling data from all animals (Figure 3.b). Average values of tissue pO_2_ increased from approximately 20.6 ± 2.9 mmHg at baseline to 28.7±2.1 mmHg at Day 3, 34.8±3.0 mmHg at Day 7). Peak values in pO₂ distributions increased from approximately 13 mmHg at baseline to 29 mmHg at Day 3, and 39.5 mmHg at Day 7.

**Figure 3.**
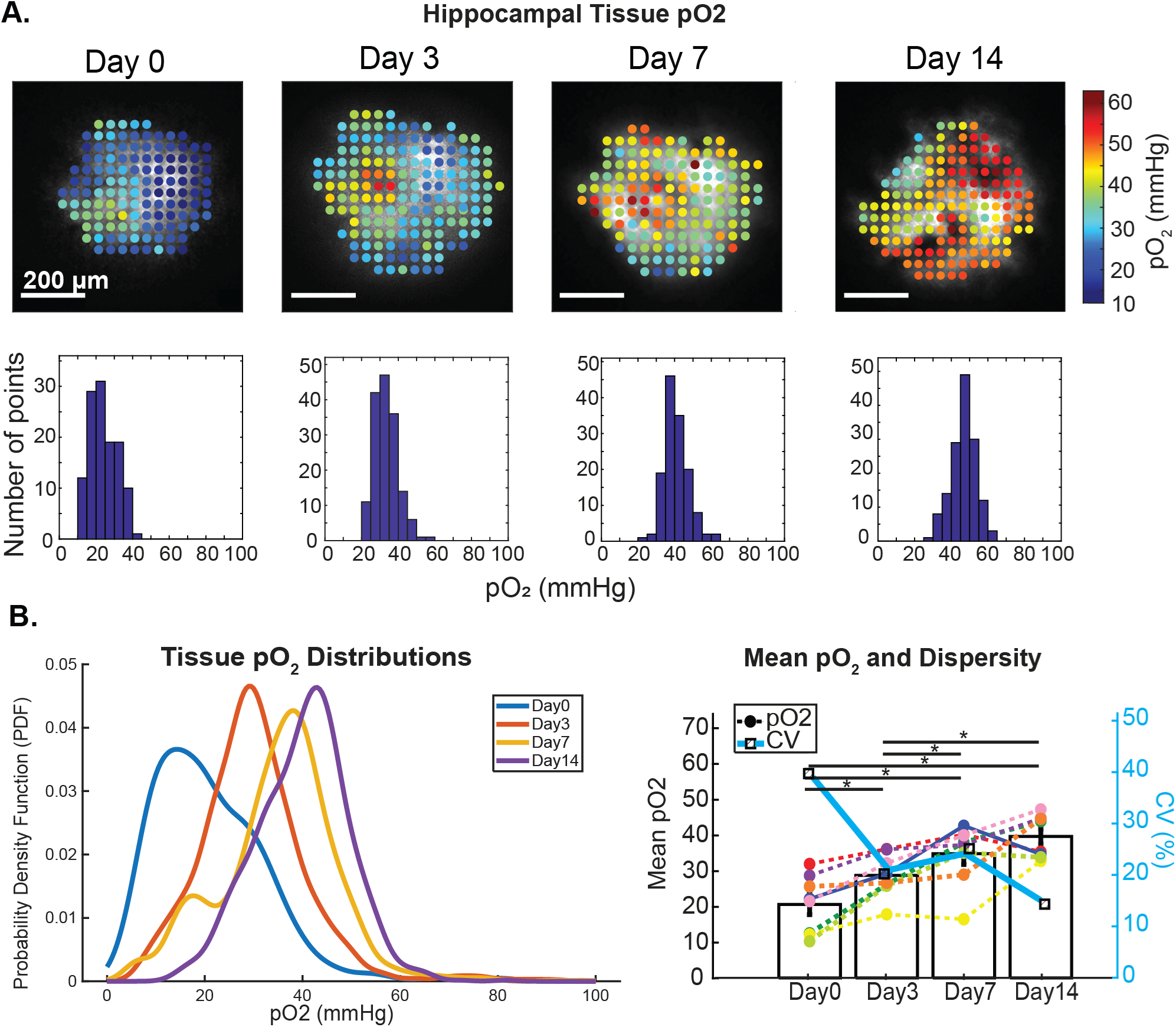
(a) Representative tissue pO₂ maps from the awake mouse CA1 region, imaged with 2PM through a GRIN lens. Top: color-coded pO₂ maps. Bottom: corresponding pO₂ distributions. (b) Pooled pO₂ distributions and dispersity from all animals. CA1 oxygen progressively increased and became less dispersed during and after partial SD (n=8)

Similar to our observations of neuronal activity, we observed that pO_2_ levels in CA1 tissue did not return to their baseline levels after restoring normal sleep and diet. After 7 days of recovery, average CA1 tissue pO₂ actually increased slightly to 39.6±2.1 mmHg while peak pO₂ increased to 40.5 mmHg. Together, the reductions in neuronal activity and increases in pO₂ indicate strongly limited benefits to recovery sleep in the HC CA1 region.

### Partial SD triggers reversible reductions to body weight

All sleep-deprived mice, with and without surgical implantation, experienced reductions in body weight during the 7-day partial SD period (Figure S4). Animals that underwent surgery, partial SD, and imaging (cohort 1) experienced average bodyweight reductions of 15.8% from Day 0 - 7. This reduction is slightly larger than the 12.2% reduction observed in the age-matched, sleep-deprived cagemates that did not receive surgery from days 0 - 7 (cohort 2). Immediately after restoring normal sleep and diet on day 8, body weight in both groups began to recover. Cohort 1 exhibited a slightly faster recovery trajectory than cohort 2. By the final day of weight collection, both groups had returned within ± 2% of their baseline weight. Cohort 3 consisted of control mice that did not undergo SD but consumed the recovery supplement for the first seven days. This group’s body weight began to decrease slightly around Day 4 and stabilized around Day 10. Although their body weights did not fully return to baseline by day 40, fluctuations were not significant (–2.2% relative to baseline).

## Discussion and Conclusion

Our longitudinal intravital imaging study revealed SD-induced reductions of spontaneous neuronal activity and corresponding increases in tissue pO_2_ within the CA1 region of mice, and a 7-day period of recovery with normal sleep and diet did not reverse the effects. In both animals and humans, numerous studies have repeatedly demonstrated that shortened sleep induces progressive reductions in vigilance, memory, and performance on cognitive and functional tasks [1,7,11]. The HC’s heightened vulnerability to SD is especially critical due to its indispensable contributions to memory formation and consolidation, navigation, learning, and other vital processes [8]. In human studies that evaluated the restorative benefits of recovery sleep, recovery periods between 7 hours to 7 days were insufficient to fully counteract the cognitive and memory declines induced by 1-10 day periods of partial SD [18,19,46,47]. Intravital visualizations of SD’s impact on cellular and molecular signaling in the live HC have only recently become possible, thanks to advances imaging advancements with GRIN lenses. Our injection-permeable cannula extends capabilities for longitudinal investigations by enabling both high-resolution imaging and repeated administration of dyes or pharmacological reagents directly to subcortical brain tissue. While rigorous comparisons between our findings and prior studies are limited by experimental differences in study design, methodologies, species, and readouts, our observations generally align well with previous reports showing SD-induced reductions in neuronal excitability, energy metabolism, and limited benefits of recovery sleep [21,48,49].

### Baseline CA1 activity

Prior to partial SD (at Day 0), we observed sparse, spontaneous burst events from CA1 Thy1 neurons, with high inter-neuron variability in both duration and frequency. Qualitatively, the sparsity and high heterogeneity are consistent with well-characterized log-normal firing rate distributions in pyramidal hippocampal neurons, observed through both *in vivo* electrophysiology and optical modalities [50–52]. Direct quantitative comparisons of metrics such as ΔF/F and firing counts from prior studies are confounded due to methodological differences, including behavioral state during recording, differing hippocampal subregions, field of view, and optical access method [43,53–56]. Nevertheless, although the fields of view, subfields, and fluorescence indicators differed between studies, our observed baseline spontanous event rate and mean duration (25 events in 8 minutes = ∼0.05 Hz, and ∼2.1 s per neuron, respectively) agree well with prior GCaMP6f recordings from awake CA1 pyramidal neurons, which report a median transient rate of 0.05 Hz [42] and transient durations on the order of 1–3 s [43]. At the network level, baseline pairwise correlation values between spontaneously active neurons were low and sparse. Numerous studies report similar observations of low overall recruitment and low correlation between pyramidal cells in CA1 and CA3 neurons during quiet wakefulness. Our findings are consistent with prior reports contending a baseline state of sparse but structured network coordination between CA1 neurons [42,43,57]. At Day 0, the PSD of φ(t) displayed local maxima around 0.18, 0.3, and 0.4 Hz, with corresponding periodicities around ∼ 5–6 seconds, ∼3s, and ∼2.5 s. Future investigations will extensively characterize the physiological bases of these peaks, including potential correlations with infraslow oscillations and sharp wave ripples. [58]

Baseline CA1 pO₂ measurements showed a mean value of 20 mmHg with relatively large heterogeneity. Our pO₂ observations agree well with recent findings from Cai et al, who performed measurements using custom-fabricated optoelectronic implants containing the PtTFPP oxygen sensor. In awake, freely-moving C57BL/6N mice, they observed tissue pO₂ values ranging between 25-30 mmHg from the HC CA3 region [59]. Notably, our tissue pO₂ measurements in the CA1 are lower than observations reported in the somatosensory cortex of awake mice. In cortical layers I-IV, Mächler et al observed mean tissue pO₂ of ∼40 mmHg with high hetereogeneity [44]. Discrepancies between our findings and Mächler et al’s may partially stem from proximity to vasculature, as their measurements specifically targeted tissue near oxygen-rich arterioles to enable calculations of cerebral metabolic rate of oxygen (CMRO2). More likely, the difference can be attributed to the HC’s lower vascularity and decreased neurovascular coupling. As indicated by simulation models, the HC’s lower capillary density and oxygen concentration result in lower tissue oxygen levels relative to the cortex [60]. Earlier studies using Clark microelectrodes and anesthetized animals reported higher tissue pO₂ in both the HC and cortex. In gerbils anesthetized with sodium pentobarbital, tissue pO₂ levels in the cortex and HC did not differ significantly (35.4 vs 33.6 mmHg) [61]. While anesthesia under isoflurane is known to elevate tissue pO₂ by 1.5–1.7 fold relative to awake states, the effects may vary across anesthetic agents and brain regions [59,62].

### Partial SD

SD’s reported effects on HC neurons vary considerably across species, SD paradigm, and measurement timepoint, hindering direct comparisons with our findings. Our observations indicate that a 7-day period of partial SD reduces the number of spontaneously-active neurons and diminishes neuronal firing rates in the awake mouse CA1. These results support prior slice electrophysiology studies, which showed that 24h of sleep fragmentation reduced excitability and long term potentiation in CA1 rat pyramidal neurons [63,64], and with reports that acute SD leads to rapid retraction of CA1 dendritic spines and impaired synaptic connectivity [65].

In apparent contrast, *in vivo* recordings during and after 9 hours acute forced SD in rats showed that pyramidal CA1 neurons maintained sustained firing during SD. In this study, Giri et al also demonstrated that sharp-wave ripples occurred at sustained or elevated rates during SD, but with reduced power and higher ripple frequency. Notably, despite the preserved firing, hippocampal replay and memory reactivation were significantly diminished, indicating that the integrity of network-level coordination is a functionally critical variable disrupted by sleep loss rather than raw firing rate alone [49]. This interpretation aligns with our own network-level observations. We observed a reduction in the fraction of co-active neurons from Day 0 to Day 3, which never fully recovered to baseline levels by Day 14.The result indicates a reduction in network co-activation, independent of the concurrent decrease in individually active neurons. Moreover, spectral analysis of the population co-activation signal φ(t) revealed a persistent reduction in broadband power from 0–1 Hz, with the loss of discrete spectral peaks at approximately 0.18, 0.3, and 0.4 Hz that were present at baseline. This indicates that partial SD disrupts not merely the rate but the rhythmic temporal organization of hippocampal population co-activation during quiet wakefulness. The diminished spectral power within φ(t) is consistent with broader evidence that sleep loss weakens hippocampal network synchrony and destabilizes coordinated population dynamics [49]

Notably, when compared with a recent two-photon Ca²⁺ imaging study, our study provides additional context regarding the impacts of sleep and sleep deprivation on HC neurons. By imaging CA1 pyramidal neurons across natural sleep-wake cycles in mice, Niethard et al demonstrated that hippocampal neuronal recruitment and Ca²⁺ signaling are higher during sleep than during quiet wakefulness. CA1 neuronal Ca^2+^ activity peaked during REM sleep and gradually reduced with subsequent slow wave sleep epochs [57]. Conversely, by curtailing sleep and extending quiet wakefulness, our study indicates that chronic partial SD reduces cumulative hippocampal population activity and disrupts rhythmically structured co-activation patterns that characterize the resting CA1 network. Taken together, Niethard et al’s and our findings highlight the CA1 region’s critical dependence on sleep and vulnerability to insufficient sleep.

In contrast to the reduction in neuronal firing counts and co-activation, pO₂ within CA1 tissue progressively increased throughout the 7-day partial SD period. Tissue pO₂ reflects the intricate balance between oxygen supply, delivered via cerebral blood flow (CBF), and oxygen demand and consumption, driven by cellular metabolism. From our study, the impact of partial SD on HC vascular oxygen supply cannot be determined due to the absence of CBF and microvascular oxygenation measurements. However, our neuronal measurements provide insight into partial SD’s impact on oxygen demand. Prolonged forced wakefulness notably elevates basal energetic costs throughout the body, increasing demand for oxygen to generate ATP. In the brain, though, because cerebral energy metabolism is predominantly driven by neuronal NA-K-ATPase pump activity to reverse ion fluxes of axon potentials and post-synaptic currents [66], SD-induced suppression of neuronal firing likely outweighs SD-related increases in basal metabolic cost [67]. We hypothesize that the net effect of partial SD is a pronounced reduction in oxygen consumption. This reduction in oxygen consumption is likely to exceed concurrent reduction in vascular oxygen supply, resulting in a net elevation of tissue pO₂. In the cortex, prior studies indicate that SD diminishes stimulus-induced functional hyperemia. SD’s effects likely also extend to other critical brain regions and may be exacerbated compared to the cortex, including the HC [68]. Using 2PM and modeling, Shaw et al reported that vascular network differences, along with pericyte and endothelial cell function, account for decreased oxygen availability and neurovascular coupling compared to neocortex. Under these conditions, the reduced neuronal activity can be reasonably interpreted as a key driver of tissue oxygen dynamics rather than a downstream consequence of altered oxygen availability. The observed elevations in tissue pO_2_ therefore reflect a state in which metabolic demand, rather than supply, is the limiting factor. Future investigations will explore this notion further by incorporating measurements of microvascular flow and oxygenation. Collectively, the HC’s spontaneous surges in neuronal metabolic demand and sub-optimal vascular supply could account for its heightened vulnerability to SD-induced alterations [8].

### Limited benefits of Recovery Sleep

After returning to normal sleep conditions from Day 8–14, neuronal firing counts exhibited only a modest, incomplete rebound by Day 14. At the network level, neither the spontaneously firing neurons nor the PSD of φ(t) recovered appreciably during this period. Conversely, tissue pO₂ continued to rise modestly from Day 7 to 14, diverging from the trajectory of neuronal recovery. Together, these findings support a model in which chronic SD selectively induces long-lasting, cumulative damage to the brain that persists beyond recovery, and metabolic restoration decouples from functional recovery.

Despite differences in experimental readouts and recovery durations, our findings align with a growing body of evidence demonstrating the limited restorative capacity of recovery sleep following sustained sleep restriction. In rats subjected to 10-day MMPM sleep restriction, a 4 h recovery period failed to restore IBA-1 immunoreactivity, synaptophysin expression, or pERK/ERK signaling to baseline levels [21]. In rats, reactivation of hippocampal neuronal firing patterns during sharp-wave ripples partially rebounded after recovery sleep but failed to reach levels observed during natural sleep [49]. A study performed by Chai et al in humans identified a dissociation between neural network normalization and behavioral recovery. After 2 nights of recovery sleep, they observed restored functional connectivity between the hippocampus and the default mode network following total SD, yet episodic memory performance remained impaired [18]. Longer recovery periods have fared little better: seven days of recovery were insufficient to reverse the effects of 10 days of 30% sleep restriction on task-evoked and resting-state EEG potentials [19]. Together, these findings compellingly demonstrate that recovery sleep cannot compensate for the cumulative burden imposed by chronic SD on hippocampal structure and function. Our study extends the notion of inadequate recovery to the metabolic function. We observed continued elevation of CA1 tissue pO₂ during recovery concurrent with restored bodyweight. When coupled with our observations of persistent suppression of neuronal firing and network co-activation, our study suggests a dissociation between systemic metabolic normalization and local hippocampal functional recovery. This dissociation suggests a post-SD transition from functionally adaptive reduction in hippocampal metabolic demand to a state of persistent metabolic-functional uncoupling during recovery. During recovery oxygen availability is no longer a limiting factor but neuronal activity nonetheless fails to normalize. This dissociation may represent a mechanistic window into how chronic partial sleep restriction elicits cumulative, lasting damage to hippocampal circuits that persists beyond the resolution of systemic sleep debt.

Our longitudinal study characterizes a previously unknown cumulative burden imposed by chronic partial SD on hippocampal neuronal signaling and tissue metabolism, revealing a persistent dissociation between metabolic and functional recovery that outlasts the sleep restriction period itself. The injection-permeable GRIN lens cannula system opens new avenues for long-term in vivo investigation of deep brain environments in preclinical models, and future experiments will incorporate concurrent measurements of CBF and microvascular oxygenation to resolve the current study’s limitation in characterizing SD-induced mismatches between metabolic supply and demand. The present study is also limited by the absence of polysomnographic verification of sleep loss during the MMPM protocol, the restriction to a single sex and adult age range, and a single recovery observation timepoint. Future work will incorporate these additional metrics to determine whether the metabolic-functional dissociation ultimately resolves or persists with prolonged recovery, and will also explore whether direct intrahippocampal delivery of compounds can counteract SD-induced suppression of neuronal excitability, including PDE4 inhibitors adenosine A1 receptor antagonists [65,69,70]

## Acknowledgments

This work was performed with generous support from the Northeastern College of Engineering and the National Institutes of Health: NIH R01AA27097, NIH R56AG058849, R21AG085655, NIH R01CA226855, NIH U01CA280849, and NIH P01CA084203. We extend our deep gratitude to Dr. Natalie Fomin-Thunemann and Dr. Anna Devor from Boston University, Dr. Yu Feng from the Icahn School of Medicine at Mount Sinai for their guidance and support with implantation and tissue injection.

Authors contributions: Z.L. designed and fabricated the cannulae, performed the surgeries and imaging experiments, analyzed and interpreted the data, and wrote the manuscript. K.M. assisted with neuronal data analysis. A.C.R. and C.L. provided technical guidance on surgical and experimental procedures and assisted with the implementation and adaptation of data analysis workflows. Y.L., P.S., and K.Z. contributed to the development and optimization of cannula preparation procedures. B.Q.S. provided guidance related to manuscript writing. M.A.Y. supervised the study, contributed to the interpretation of the results and writing of the manuscript, and reviewed and edited the manuscript. The authors declare that they have no competing interests. The data and code supporting the findings of this study will be made publicly available upon publication. Until then, they are available from the corresponding author upon reasonable request. AI tools were used to assist in the initial drafting of the analysis code. All AI-assisted code was subsequently reviewed, modified, and validated by the author, who takes full responsibility for the final code and analyses.

## References

1 R. F. Gottesman, P. L. Lutsey, H. Benveniste, D. L. Brown, K. M. Full, J.-M. Lee, R. S. Osorio, M. P. Pase, N. S. Redeker, S. Redline, A. P. Spira, and on behalf of the American Heart Association Stroke Council; Council on Cardiovascular and Stroke Nursing; and Council on Hypertension, “Impact of Sleep Disorders and Disturbed Sleep on Brain Health: A Scientific Statement From the American Heart Association,” Stroke 55, (2024).

2. H. R. Colten, B. M. Altevogt, and I. of M. (US) C. on S. M. and Research, “Extent and Health Consequences of Chronic Sleep Loss and Sleep Disorders,” in Sleep Disorders and Sleep Deprivation: An Unmet Public Health Problem (National Academies Press (US), 2006).

3. V. K. Chattu, Md. D. Manzar, S. Kumary, D. Burman, D. W. Spence, and S. R. Pandi-Perumal, “The Global Problem of Insufficient Sleep and Its Serious Public Health Implications,” Healthcare 7, 1 (2018).

4. D. Kocevska, T. S. Lysen, A. Dotinga, M. E. Koopman-Verhoeff, M. P. C. M. Luijk, N. Antypa, N. R. Biermasz, A. Blokstra, J. Brug, W. J. Burk, H. C. Comijs, E. Corpeleijn, H. S. Dashti, E. J. de Bruin, R. de Graaf, I. P. M. Derks, J. F. Dewald-Kaufmann, P. J. M. Elders, R. J. B. J. Gemke, L. Grievink, L. Hale, C. A. Hartman, C. J. Heijnen, M. Huisman, A. Huss, M. A. Ikram, S. E. Jones, M. K. Velderman, M. Koning, A. M. Meijer, K. Meijer, R. Noordam, A. J. Oldehinkel, J. O. Groeniger, B. W. J. H. Penninx, H. S. J. Picavet, S. Pieters, S. A. Reijneveld, E. Reitz, C. M. Renders, G. Rodenburg, F. Rutters, M. C. Smith, A. S. Singh, M. B. Snijder, K. Stronks, M. ten Have, J. W. R. Twisk, D. Van de Mheen, J. van der Ende, K. B. van der Heijden, P. G. van der Velden, F. J. van Lenthe, R. R. L. van Litsenburg, S. H. van Oostrom, F. J. van Schalkwijk, C. M. Sheehan, R. A. Verheij, F. C. Verhulst, M. C. M. Vermeulen, R. C. H. Vermeulen, W. M. M. Verschuren, T. G. M. Vrijkotte, A. H. Wijga, A. M. Willemen, M. ter Wolbeek, A. R. Wood, Y. Xerxa, W. M. Bramer, O. H. Franco, A. I. Luik, E. J. W. Van Someren, and H. Tiemeier, “Sleep characteristics across the lifespan in 1.1 million people from the Netherlands, United Kingdom and United States: a systematic review and meta-analysis,” Nat. Hum. Behav. 5, 113–122 (2021).

5. A. V. Benjafield, F. H. Sert Kuniyoshi, A. Malhotra, J. L. Martin, C. M. Morin, L. F. Maurer, P. A. Cistulli, J.-L. Pépin, and E. M. Wickwire, “Estimation of the global prevalence and burden of insomnia: a systematic literature review-based analysis,” Sleep Med. Rev. 82, 102121 (2025).

6. L. Anghel, A. Ciubară, A. Nechita, L. Nechita, C. Manole, L. Baroiu, A. B. Ciubară, and C. L. Mușat, “Sleep Disorders Associated with Neurodegenerative Diseases,” Diagnostics 13, 2898 (2023).

7. Z. Zamore and S. C. Veasey, “Neural consequences of chronic sleep disruption,” Trends Neurosci. 45, 678–691 (2022).

8. R. Havekes and T. Abel, “The tired hippocampus: the molecular impact of sleep deprivation on hippocampal function,” Curr. Opin. Neurobiol. 44, 13–19 (2017).

9. J. Feng, X. Zhao, G. G. Gurkoff, K. C. Van, K. Shahlaie, and B. G. Lyeth, “Post-Traumatic Hypoxia Exacerbates Neuronal Cell Death in the Hippocampus,” J. Neurotrauma 29, 1167–1179 (2012).

10. P. Malerba and M. Bazhenov, “Circuit mechanisms of hippocampal reactivation during sleep,” Neurobiol. Learn. Mem. 160, 98–107 (2019).

11. S. Diekelmann, I. Wilhelm, and J. Born, “The whats and whens of sleep-dependent memory consolidation,” Sleep Med. Rev. 13, 309–321 (2009).

12. K. Mizuseki and H. Miyawaki, “Hippocampal information processing across sleep/wake cycles,” Neurosci. Res. 118, 30–47 (2017).

13. D. Koukaroudi, Z. Qiu, E. Fransén, R. Gokhale, E. Bulovaite, N. H. Komiyama, J. Seibt, and S. G. N. Grant, “Sleep maintains excitatory synapse diversity in the cortex and hippocampus,” Curr. Biol. 34, 3836–3843.e5 (2024).

14. Z. Guan, X. Peng, and J. Fang, “Sleep deprivation impairs spatial memory and decreases extracellular signal-regulated kinase phosphorylation in the hippocampus,” Brain Res. 1018, 38–47 (2004).

15. R. Zhao, X. Zhang, Y. Zhu, N. Fei, J. Sun, P. Liu, X. Yang, and W. Qin, “Disrupted Resting-State Functional Connectivity in Hippocampal Subregions After Sleep Deprivation,” Neuroscience 398, 37–54 (2019).

16. J. Leerssen, R. Wassing, J. R. Ramautar, D. Stoffers, O. Lakbila-Kamal, J. Perrier, J. Bruijel, J. C. Foster-Dingley, M. Aghajani, and E. J. W. van Someren, “Increased hippocampal-prefrontal functional connectivity in insomnia,” Neurobiol. Learn. Mem. 160, 144–150 (2019).

17. S. P. Feeney, J. M. McCarthy, C. R. Petruconis, and J. C. Tudor, “Sleep loss is a metabolic disorder,” Sci Signal (2025).

18. Y. Chai, Z. Fang, F. N. Yang, S. Xu, Y. Deng, A. Raine, J. Wang, M. Yu, M. Basner, N. Goel, J. J. Kim, D. A. Wolk, J. A. Detre, D. F. Dinges, and H. Rao, “Two nights of recovery sleep restores hippocampal connectivity but not episodic memory after total sleep deprivation,” Sci. Rep. 10, 8774 (2020).

19. J. K. Ochab, J. Szwed, K. Oleś, A. Bereś, D. R. Chialvo, A. Domagalik, M. Fąfrowicz, H. Ogińska, E. Gudowska-Nowak, T. Marek, and M. A. Nowak, “Observing changes in human functioning during induced sleep deficiency and recovery periods,” PLOS ONE 16, e0255771 (2021).

20. B. Giri, N. Kinsky, U. Kaya, K. Maboudi, T. Abel, and K. Diba, “Sleep loss diminishes hippocampal reactivation and replay,” Nature 630, 935–942 (2024).

21. J. E. García-Aviles, J. J. Avilez-Avilez, J. Sánchez-Hernández, C. Patlán-Márquez, J. Rodríguez-Alpízar, F. M. Becerril-Mercado, A. Jiménez, N. N. Guerrero-Vargas, J.-P. Morin, M. Rodríguez-García, J. Manjarrez-Marmolejo, B. Gómez-González, R. Guevara-Guzmán, and M. A. Guzmán-Ruiz, “A Sleeping Opportunity Does Not Restore Hippocampal Alterations Induced by 10 Days of Sleep Restriction in Rats,” Neurochem. Res. 50, 311 (2025).

22. V. Colavito, P. F. Fabene, G. Grassi-Zucconi, F. Pifferi, Y. Lamberty, M. Bentivoglio, and G. Bertini, “Experimental sleep deprivation as a tool to test memory deficits in rodents,” Front. Syst. Neurosci. 7, (2013).

23. M. E. Bocarsly, W. Jiang, C. Wang, J. T. Dudman, N. Ji, and Y. Aponte, “Minimally invasive microendoscopy system for in vivo functional imaging of deep nuclei in the mouse brain,” Biomed. Opt. Express 6, 4546 (2015).

24. H. Salmon, M. R. Rasouli, N. Distasio, and M. Tabrizian, “Facile engineering and interfacing of styrenic block copolymers devices for low-cost, multipurpose microfluidic applications,” Eng. Rep. 3, e12361 (2021).

25. M. Etayo-Escanilla, N. Campillo, P. Ávila-Fernández, J. M. Baena, J. Chato-Astrain, F. Campos, D. Sánchez-Porras, Ó. D. García-García, and V. Carriel, “Comparison of Printable Biomaterials for Use in Neural Tissue Engineering: An In Vitro Characterization and In Vivo Biocompatibility Assessment,” Polymers 16, 1426 (2024).

26. H. Dana, T.-W. Chen, A. Hu, B. C. Shields, C. Guo, L. L. Looger, D. S. Kim, and K. Svoboda, “Thy1-GCaMP6 Transgenic Mice for Neuronal Population Imaging In Vivo,” PLoS ONE 9, e108697 (2014).

27. S. L. Resendez, J. H. Jennings, R. L. Ung, V. M. K. Namboodiri, Z. C. Zhou, J. M. Otis, H. Nomura, J. A. McHenry, O. Kosyk, and G. D. Stuber, “Visualization of cortical, subcortical and deep brain neural circuit dynamics during naturalistic mammalian behavior with head-mounted microscopes and chronically implanted lenses,” Nat. Protoc. 11, 566–597 (2016).

28. V. C. Rossi, P. A. Tiba, K. D. M. Moreira, T. L. Ferreira, M. G. M. Oliveira, and D. Suchecki, “Effects of sleep deprivation on different phases of memory in the rat: dissociation between contextual and tone fear conditioning tasks,” Front. Behav. Neurosci. 8, (2014).

29. M. Mônico-Neto, S. Q. D. C. Giampá, K. S. Lee, C. M. De Melo, H. D. S. Souza, M. Dáttilo, P. A. Minali, P. H. Santos Prado, S. Tufik, M. T. De Mello, and H. K. M. Antunes, “Negative Energy Balance Induced by Paradoxical Sleep Deprivation Causes Multicompartmental Changes in Adipose Tissue and Skeletal Muscle,” Int. J. Endocrinol. 2015, 1–6 (2015).

30. C. Wang, W.-R. Gao, J. Yin, Z.-J. Wang, J.-S. Qi, H.-Y. Cai, and M.-N. Wu, “Chronic sleep deprivation exacerbates cognitive and synaptic plasticity impairments in APP/PS1 transgenic mice,” Behav. Brain Res. 412, 113400 (2021).

31. H. Diao, Y. Li, W. Sun, J. Zhang, M. Wang, Y. Chen, F. Zhou, and X. Li, “REM sleep deprivation induced by the modified multi-platform method has detrimental effects on memory: A systematic review and meta-analysis,” Behav. Brain Res. 454, 114652 (2023).

32. T. V. Esipova, M. J. P. Barrett, E. Erlebach, A. E. Masunov, B. Weber, and S. A. Vinogradov, “Oxyphor 2P: A High-Performance Probe for Deep-Tissue Longitudinal Oxygen Imaging,” Cell Metab. 29, 736–744.e7 (2019).

33. A. Giovannucci, J. Friedrich, P. Gunn, J. Kalfon, B. L. Brown, S. A. Koay, J. Taxidis, F. Najafi, J. L. Gauthier, P. Zhou, B. S. Khakh, D. W. Tank, D. B. Chklovskii, and E. A. Pnevmatikakis, “CaImAn an open source tool for scalable calcium imaging data analysis,” eLife 8, e38173 (2019).

34. E. A. Pnevmatikakis, D. Soudry, Y. Gao, T. A. Machado, J. Merel, D. Pfau, T. Reardon, Y. Mu, C. Lacefield, W. Yang, M. Ahrens, R. Bruno, T. M. Jessell, D. S. Peterka, R. Yuste, and L. Paninski, “Simultaneous Denoising, Deconvolution, and Demixing of Calcium Imaging Data,“ Neuron 89, 285–299 (2016).

35. L. Reznichenko, Q. Cheng, K. Nizar, S. L. Gratiy, P. A. Saisan, E. M. Rockenstein, T. Gonzalez, C. Patrick, B. Spencer, P. Desplats, A. M. Dale, A. Devor, and E. Masliah, “In Vivo Alterations in Calcium Buffering Capacity in Transgenic Mouse Model of Synucleinopathy,” J. Neurosci. 32, 9992–9998 (2012).

36. J. Graf, V. Rahmati, M. Majoros, O. W. Witte, C. Geis, S. J. Kiebel, K. Holthoff, and K. Kirmse, “Network instability dynamics drive a transient bursting period in the developing hippocampus in vivo,” eLife 11, e82756 (2022).

37. T. Sasaki, T. Ishikawa, R. Abe, R. Nakayama, A. Asada, N. Matsuki, and Y. Ikegaya, “Astrocyte calcium signalling orchestrates neuronal synchronization in organotypic hippocampal slices,” J. Physiol. 592, 2771–2783 (2014).

38. P. R. Bressan, E. Long, J. Jiang, R. Vithayathil, Z. Guan, Y. Song, B. C. Rauscher, N. Chai, K. Kılıç, Ş. E. Erdener, A. Devor, D. A. Boas, and R. P. Tang, “Visual gamma stimulation causes prolonged enhancement of low-frequency blood flow oscillations across cortical regions in mice,” 2026.05.31.729102 (2026).

39. V. Saravanan, G. J. Berman, and S. J. Sober, “Application of the Hierarchical Bootstrap to Multi-Level Data in Neuroscience,” (2019).

40. P. Ranganathan, “An Introduction to Statistics: Choosing the Correct Statistical Test,” Indian J. Crit. Care Med. 25, S184–S186 (2021).

41. V. Saravanan, G. J. Berman, and S. J. Sober, “Application of the Hierarchical Bootstrap to Multi-Level Data in Neuroscience,” 819334 (2020).

42. V. Villette, M. Levesque, A. Miled, B. Gosselin, and L. Topolnik, “Simple platform for chronic imaging of hippocampal activity during spontaneous behaviour in an awake mouse,” Sci. Rep. 7, 43388 (2017).

43. G. Schoenfeld, S. Carta, P. Rupprecht, A. Ayaz, and F. Helmchen, “In Vivo Calcium Imaging of CA3 Pyramidal Neuron Populations in Adult Mouse Hippocampus,” eNeuro 8, (2021).

44. P. Mächler, N. Fomin-Thunemann, M. Thunemann, M. J. Sætra, M. Desjardins, K. Kılıç, L. N. Amra, E. A. Martin, I. A. Chen, I. Şencan-Eğilmez, B. Li, P. Saisan, J. X. Jiang, Q. Cheng, K. L. Weldy, D. A. Boas, R. B. Buxton, G. T. Einevoll, A. M. Dale, S. Sakadžić, and A. Devor, “Baseline oxygen consumption decreases with cortical depth,” PLOS Biol. 20, e3001440 (2022).

45. M. Rakrak, “Central tendency and data distribution: How to choose the right measure for accurate analysis,” Int. J. Lit. Educ. 5, 71–75 (2025).

46. J. Axelsson, G. Kecklund, T. Åkerstedt, P. Donofrio, M. Lekander, and M. Ingre, “Sleepiness and Performance in Response to Repeated Sleep Restriction and Subsequent Recovery during Semi-Laboratory Conditions,” Chronobiol. Int. 25, 297–308 (2008).

47. K. Ikegami, S. Ogyu, Y. Arakomo, K. Suzuki, K. Mafune, H. Hiro, and S. Nagata, “Recovery of Cognitive Performance and Fatigue after One Night of Sleep Deprivation,” J. Occup. Health 51, 412–422 (2009).

48. C. M. McDermott, G. J. LaHoste, C. Chen, A. Musto, N. G. Bazan, and J. C. Magee, “Sleep Deprivation Causes Behavioral, Synaptic, and Membrane Excitability Alterations in Hippocampal Neurons,“ J. Neurosci. 23, 9687–9695 (2003).

49. B. Giri, N. Kinsky, U. Kaya, K. Maboudi, T. Abel, and K. Diba, “Sleep loss diminishes hippocampal reactivation and replay,” Nature 630, 935–942 (2024).

50. K. Mizuseki and G. Buzsáki, “Preconfigured, Skewed Distribution of Firing Rates in the Hippocampus and Entorhinal Cortex,“ Cell Rep. 4, 1010–1021 (2013).

51. G. Buzsáki and K. Mizuseki, “The log-dynamic brain: how skewed distributions affect network operations,” Nat. Rev. Neurosci. 15, 264–278 (2014).

52. Y. Hayashi, K. Kobayakawa, and R. Kobayakawa, “The temporal and contextual stability of activity levels in hippocampal CA1 cells,” Proc. Natl. Acad. Sci. 120, e2221141120 (2023).

53. D. A. Dombeck, C. D. Harvey, L. Tian, L. L. Looger, and D. W. Tank, “Functional imaging of hippocampal place cells at cellular resolution during virtual navigation,” Nat. Neurosci. 13, 1433–1440 (2010).

54. M. Kondo, K. Kobayashi, M. Ohkura, J. Nakai, and M. Matsuzaki, “Two-photon calcium imaging of the medial prefrontal cortex and hippocampus without cortical invasion,” eLife 6, e26839 (2017).

55. A. Formozov, M. Chini, A. Dieter, W. Yang, J. A. Pöpplau, I. L. Hanganu-Opatz, and J. S. Wiegert, “Calcium Imaging and Electrophysiology of hippocampal Activity under Anesthesia and natural Sleep in Mice,” Sci. Data 9, 113 (2022).

56. W. Zong, H. A. Obenhaus, E. R. Skytøen, H. Eneqvist, N. L. de Jong, R. Vale, M. R. Jorge, M.-B. Moser, and E. I. Moser, “Large-scale two-photon calcium imaging in freely moving mice,” Cell 185, 1240–1256.e30 (2022).

57. N. Niethard, D. M. Pagano, N. Mojtahedi, Y. Kovalchuk, J. Born, and O. Garaschuk, “Sleep spindles promote hippocampal network downregulation during sleep,” Curr. Biol. 36, 1385–1398.e3 (2026).

58. G. Buzsáki, “Hippocampal sharp wave-ripple: A cognitive biomarker for episodic memory and planning,” Hippocampus 25, 1073–1188 (2015).

59. X. Cai, H. Zhang, P. Wei, Q. Liu, D. Sheng, Z. Li, W. Zhao, Z. Ye, Z. Xue, Y. Xie, Y. Dai, C. Wang, Y. Wang, X. Fu, B. Zhang, L. Yin, H. Peng, H. Ding, G. Zhao, and X. Sheng, “A Wireless Optoelectronic Probe Monitors Tissue Oxygenation in the Deep Brain,” 2023.05.21.541646 (2023).

60. K. Shaw, L. Bell, K. Boyd, D. M. Grijseels, D. Clarke, O. Bonnar, H. S. Crombag, and C. N. Hall, “Neurovascular coupling and oxygenation are decreased in hippocampus compared to neocortex because of microvascular differences,” Nat. Commun. 12, 3190 (2021).

61. P. K. Nair, D. G. Buerk, and J. H. Halsey, “Comparisons of oxygen metabolism and tissue PO2 in cortex and hippocampus of gerbil brain.,” Stroke 18, 616–622 (1987).

62. D. G. Lyons, A. Parpaleix, M. Roche, and S. Charpak, “Mapping oxygen concentration in the awake mouse brain,” eLife 5, e12024 (2016).

63. J. L. Tartar, J. T. McKenna, C. P. Ward, R. W. McCarley, R. E. Strecker, and R. E. Brown, “Sleep fragmentation reduces hippocampal CA1 pyramidal cell excitability and response to adenosine,” Neurosci. Lett. 469, 1–5 (2010).

64. C. M. McDermott, G. J. LaHoste, C. Chen, A. Musto, N. G. Bazan, and J. C. Magee, “Sleep Deprivation Causes Behavioral, Synaptic, and Membrane Excitability Alterations in Hippocampal Neurons,“ J. Neurosci. 23, 9687–9695 (2003).

65. R. Havekes, A. J. Park, J. C. Tudor, V. G. Luczak, R. T. Hansen, S. L. Ferri, V. M. Bruinenberg, S. G. Poplawski, J. P. Day, S. J. Aton, K. Radwańska, P. Meerlo, M. D. Houslay, G. S. Baillie, and T. Abel, “Sleep deprivation causes memory deficits by negatively impacting neuronal connectivity in hippocampal area CA1,” eLife 5, e13424 (2016).

66. J. J. Harris, R. Jolivet, and D. Attwell, “Synaptic Energy Use and Supply,” Neuron 75, 762–777 (2012).

67. D. Attwell and C. Iadecola, “The neural basis of functional brain imaging signals,” Trends Neurosci. 25, 621–625 (2002).

68. T. Csipo, A. Lipecz, C. Owens, P. Mukli, J. W. Perry, S. Tarantini, P. Balasubramanian, Á. Nyúl-Tóth, V. Yabluchanska, F. A. Sorond, J. M. Kellawan, G. Purebl, W. E. Sonntag, A. Csiszar, Z. Ungvari, and A. Yabluchanskiy, “Sleep deprivation impairs cognitive performance, alters task-associated cerebral blood flow and decreases cortical neurovascular coupling-related hemodynamic responses,” Sci. Rep. 11, 20994 (2021).

69. C. G. Vecsey, G. S. Baillie, D. Jaganath, R. Havekes, A. Daniels, M. Wimmer, T. Huang, K. M. Brown, X.-Y. Li, G. Descalzi, S. S. Kim, T. Chen, Y.-Z. Shang, M. Zhuo, M. D. Houslay, and T. Abel, “Sleep deprivation impairs cAMP signaling in the hippocampus,” Nature 461, 1122–1125 (2009).

70. C. Florian, C. G. Vecsey, M. M. Halassa, P. G. Haydon, and T. Abel, “Astrocyte-Derived Adenosine and A1 Receptor Activity Contribute to Sleep Loss-Induced Deficits in Hippocampal Synaptic Plasticity and Memory in Mice,” J. Neurosci. 31, 6956–6962 (2011).

