## Supplementary Materials for "Partial sleep deprivation diminishes and uncouples neuronal activity and energy metabolism in the live mouse hippocampus"

**Supplementary Methods**

Cannula design and fabrication

For cannula fabrication, the transparent Flexdym window material (thickness: 250 μm) was cut into circular 1mm-diameter discs using a 1mm-diameter tissue biopsy punch and a rubber mallet. The Flexdym disc was then carefully placed inside the polyimide tube at one end. A 1mm flat-end dowel pin (McMaster-Carr) was inserted into the tube to adjust the position of the disc. To enhance adhesion of both the tube and the Flexdym disc, the end of the tube with Flexdym disc was treated using a corona gun (Electro-Technic Products, model: BD-20A) for approximately two minutes at a distance of 1cm. Immediately after corona treatment, we used applied transparent optical adhesive (Norland Products, model: NOA 61) to the gap between the Flexdym disc and the tube edge using a 31G insulin needle. Then the cannula (“cannula” refers to the whole assembly after the Flexdym disc is attached to the tube) was placed vertically with the adhesive end facing upward and a custom 3D-printed stand was used to hold it upright. The cannula was exposed to a 100W 365nm UV curing lamp at a distance of 6 inches for 6-7 minutes. After the curing, the cannula was trimmed into around 4 mm in length under a microscope using dissecting scissors. The cut was made as perpendicular to the tube axis as possible. The length of the trimmed tube was measured with a caliper and the measurement was recorded. Gloves and tweezers should be used throughout the process to prevent fingerprints on Flexdym. Any surface or object that may come into contact with the Flexdym should be cleaned with water and dried with Kimwipes to prevent Flexdym from picking up dirt.

Bonding testing

A bonding test was performed to examine whether the Flexdym disc would remain firmly adhered to the polyimide tube during injection. The test was designed to mimic the intracerebral injection process. We mounted a 34G 0.375-inch needle-fitted Hamilton syringe (Hamilton) onto a microinjector (Harvard Apparatus) which controls the injection parameters, and the microinjector was secured onto a motorized micromanipulator (Scientifica) which allows precise control of the needle movement. This setup is identical to that used in the subsequent in vivo injection. The lid of a small plastic container was used as a surrogate for the skull. A hole was drilled through the lid and the cannula was fixed in place. The lid itself was held securely using adjustable mechanical fasteners, allowing stable mounting and repeatable positioning throughout the testing. The Flexdym end of the cannula was exposed to air, allowing direct observation of the Flexdym disc throughout the injection process (Figure S1.c). In real injections, the presence of brain tissue beneath the disc is expected to provide additional support, making displacement even less likely. A portable digital microscope was used to align the needle tip with the upper edge of the cannula prior to injection and to observe the needle as it passed through the Flexdym disc. During each trial, the needle was aligned to the center of the Flexdym window and advanced at a constant speed of 5 µm/s until it pierced through the disc. Six cannulae were tested in triplicate. The plastic lid was removed and repositioned between trials for re-alignment of each independent penetration. Because the Flexdym membrane is deformable, the required displacement exceeded the physical thickness of the window. For each penetration, the needle travel distance required to fully pierce the Flexdym disc was recorded. These measurements establish the needle advancement needed to reliably puncture the membrane in subsequent applications.

Injection and imaging feasibility testing

This testing consisted of two parts. Part 1 aimed to determine parameters required for a reliable injection through the Flexdym disc, including depth, speed, and volume. Part 2 evaluated whether imaging remained feasible through the punctured membrane after needle penetration. In both parts, we used the same device setup and mounted the cannula in the same way as in the bonding test. However, in contrast to the bonding test where the lid was open to air, the lid here was attached to a matching container filled with a gelatin phantom. In this configuration, the lower end of the cannula extended into the phantom, allowing needle penetration and imaging to be performed under tissue-like conditions. As a preliminary visualization test, we first performed a pilot injection using trypan blue. The phantom was prepared by dissolving 1.5 g of gelatin powder in 100 mL of a milk–deionized water mixture (1:4, v/v). The addition of milk increased the visual contrast of the injected dye, allowing the injection and resulting dye bolus to be readily observed. A total of 2000 nL of trypan blue solution was injected at 10 nL/s, and the resulting dye distribution was visually assessed (Figure S1.d). Following this pilot test, a more refined injection experiment was performed using fluorescence imaging. The gelatin phantom was prepared by dissolving 1.5 g of gelatin powder in 100 mL of deionized water, and 50 mg/mL FITC was injected into the gelatin phantom. The visible fluorescence allowed us to adjust the injection depth and volume such that the injected dye would fill the entire field of view (FOV) beneath the cannula window and, after accounting for alignment and positioning tolerances, be delivered into the phantom immediately beneath the cannula window while remaining outside the cannula lumen. The injection speed was incrementally increased from 2 nL/s to identify a rate that reliably produced a dye bolus without incomplete delivery. To further assess imaging feasibility, 5 µL of 1 µm fluorescent beads (MilliporeSigma) was diluted in 3 mL of deionized water and mixed with the gelatin solution at a 3:1 volume ratio, yielding a bead density suitable for fluorescence imaging through the punctured membrane. Images were acquired before and after injection. Fluorescence intensity profiles of individual beads were analyzed to assess changes in imaging performance, where the most closely matched Z-planes of the same beads were selected for analysis.

**Supplementary Results**

Cannula design and validation

Among the six cannulae tested in triplicate, all six Flexdym windows remained firmly bonded to the polyimide tube throughout all penetration trials, indicating adequate bonding strength. Advancing the needle approximately 1500 µm beyond the bottom of the cannula reliably perforated the flexible Flexdym membrane. Subsequent retraction of the needle by approximately 900 µm consistently enabled dye injection into the intended target region beneath the cannula window. Although the nominal needle bevel length is 750 µm, the selected advance and retraction distances (1500 µm and 900 µm) intentionally incorporated additional margins to accommodate alignment tolerances and operator-dependent calibration variability, and ensured consistent sub-window injection in practice. An injection volume of 120 nL delivered at a rate of 10 nL/s was identified as suitable for subsequent experiments. This parameter combination balanced the need to reduce the potential adverse effects associated with rapid fluid influx into brain tissue, while helping to mitigate the risk of needle clogging. In the subsequent imaging feasibility testing, between the pre- and post-injection images, slight lateral shift and axial mismatch in imaging depth were observed, both due to the re-insertion of the GRIN lens. Nevertheless, most previously imaged beads as well as the injected FITC were clearly detectable within the FOV after injection, indicating that imaging of the same region remained feasible after injection. The fluorescence intensity profiles were comparable before and after injection (FWHM= 2.62 and 3.37 µm), suggesting that the injection did not affect imaging performance (Figure S2).

**Supplementary Figures**

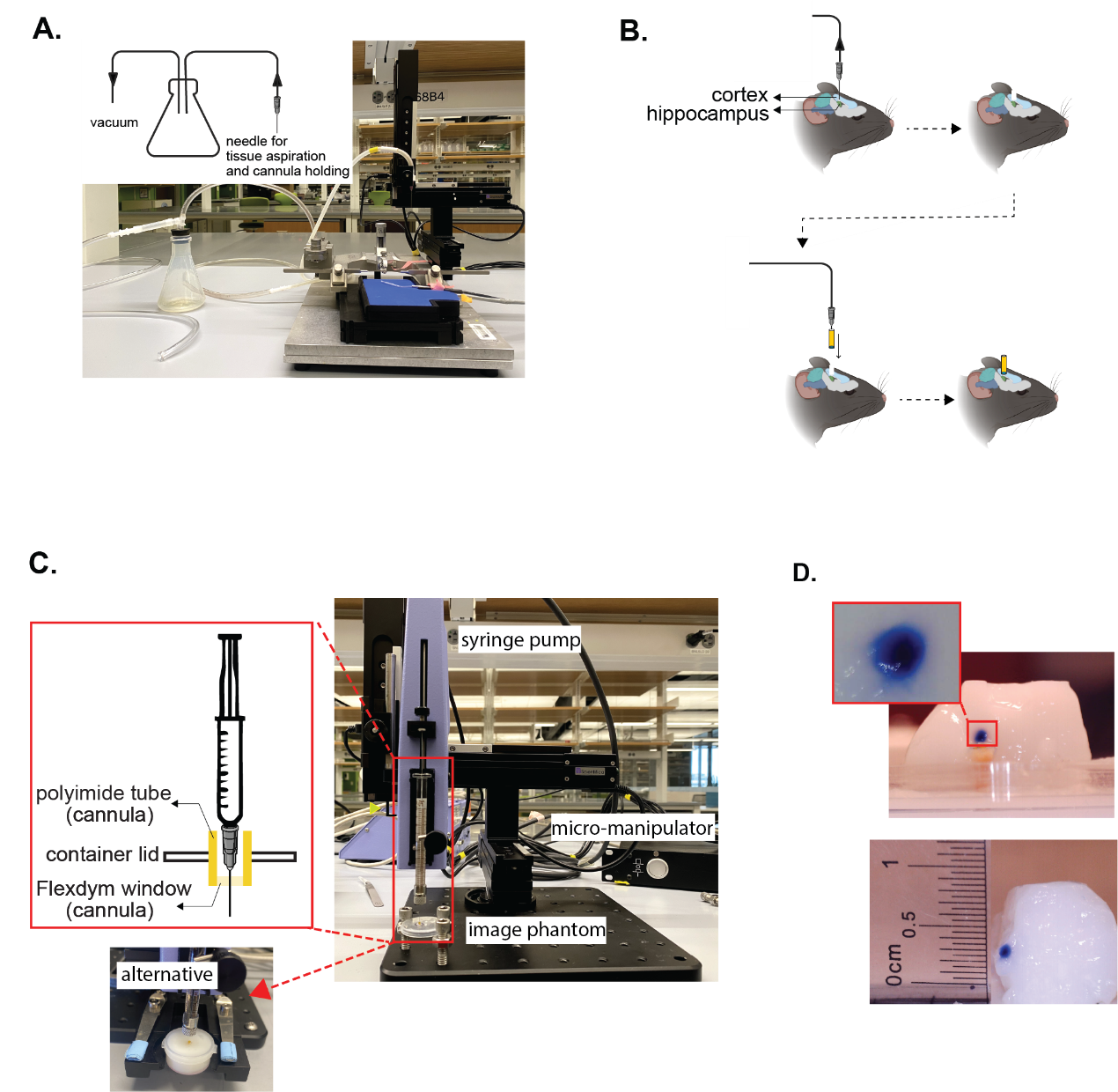

**Figure S1.** (a) Custom suction system. A blunt needle was connected to a vacuum source through a flask and mounted on a micromanipulator for controlled tissue aspiration and cannula holding. (b) Schematic of the cannula implantation procedure. Cortical tissue above the hippocampus was first aspirated using the suction system shown in (a). The same vacuum-mediated holder was then used to hold and lower the cannula into the brain, after which the cannula was secured to the skull with adhesive. Mouse illustrations were created using assets from BioRender.com. (c) Experimental setup used for bonding and injection feasibility testing. A Hamilton syringe was mounted on a syringe pump, which was secured to a motorized micromanipulator for controlled needle advancement. The cannula was fixed to a container lid with the Flexdym window either exposed to air for bonding testing or positioned in a gelatin phantom for injection and imaging feasibility testing. (d) Trypan blue distribution following pilot injection into the milk–gelatin phantom. Injection of 2000 nL at 10 nL/s resulted in an approximately 1 mm³ spherical distribution after ~2 h of diffusion.

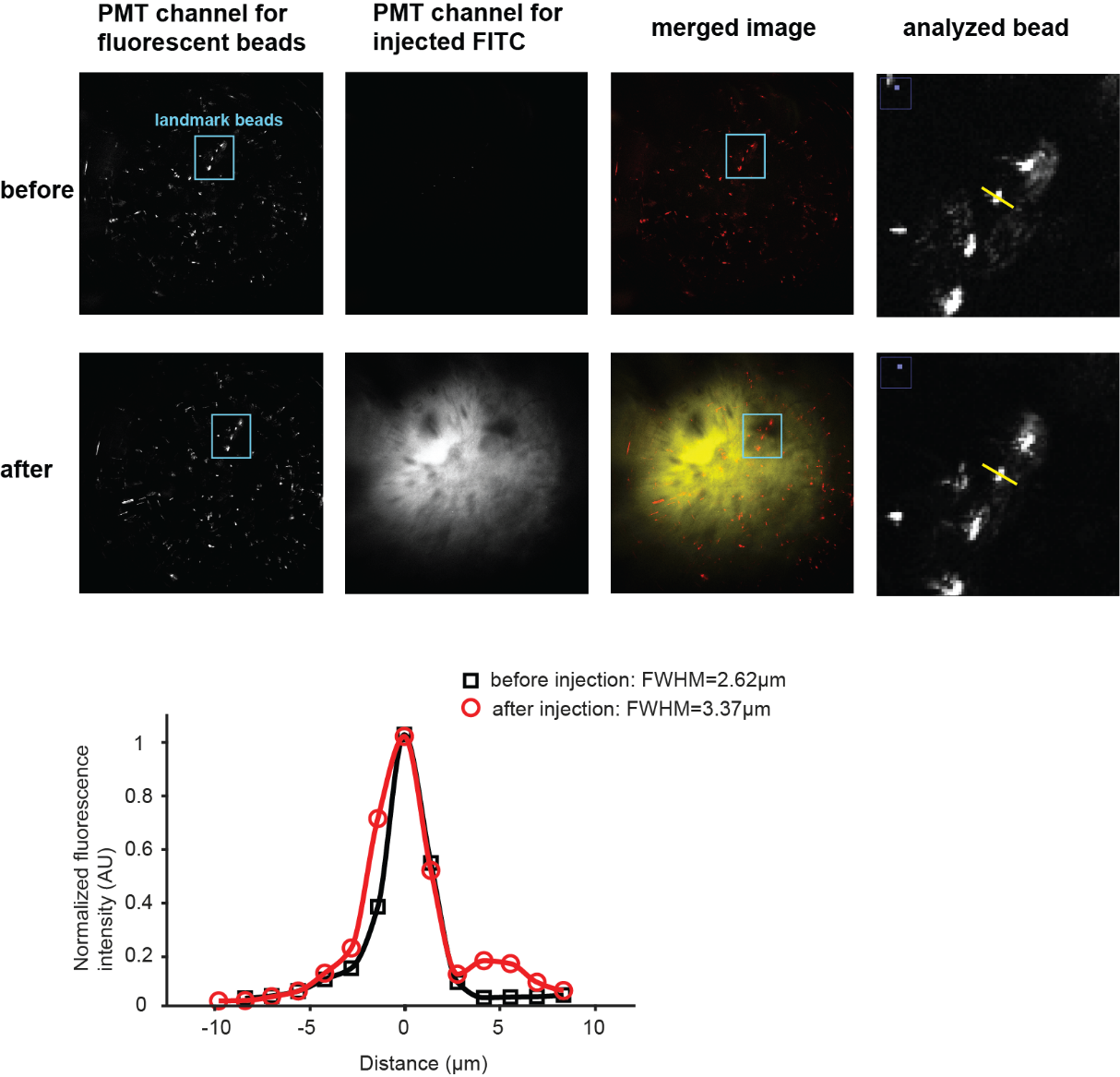

**Figure S2.** Imaging feasibility before and after injection through the Flexdym window. Representative images acquired before (first row) and after (second row) injection show fluorescent beads, injected FITC, and their merged signals. Blue boxes indicate the same landmark beads identified before and after injection. Yellow lines are used to obtain the fluorescence intensity profiles of the analyzed bead. Smooth curves were generated by interpolation of the measured intensity values for visualization. FWHM was estimated by Gaussian fitting of the primary intensity peak.

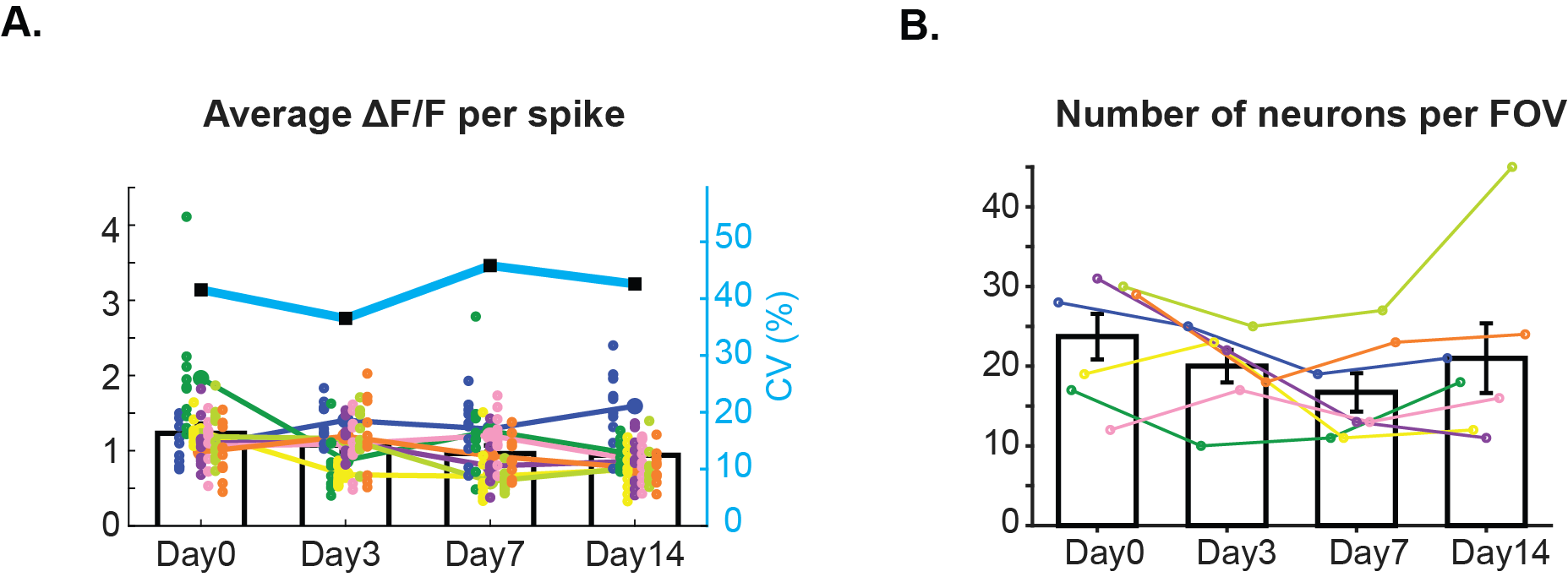

**Figure S3.** (a) Spontaneous neuronal firing amplitude and (b) Number of spontaneously active neurons, quantified before, during, and after partial SD.

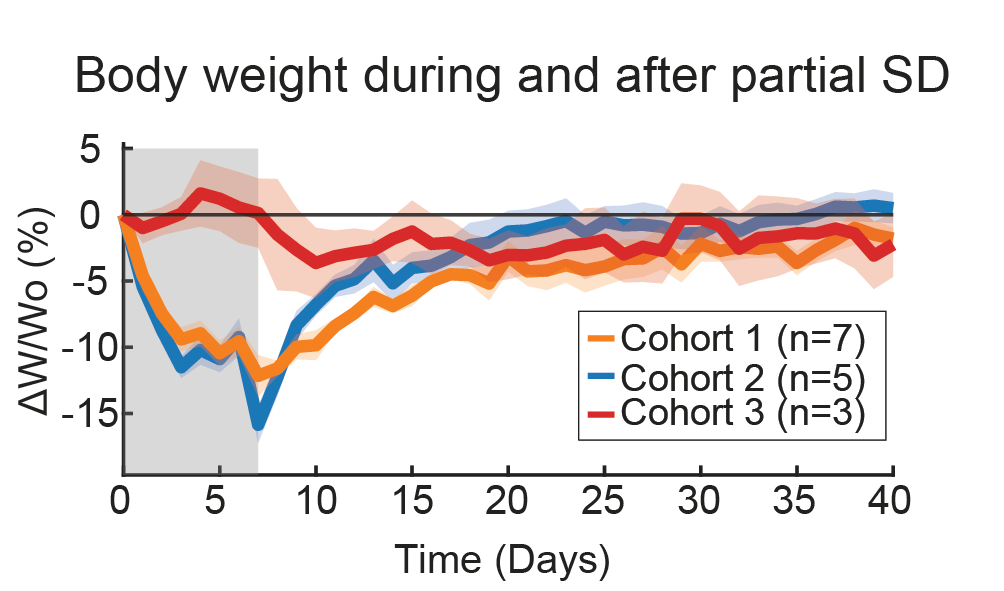

**Figure S4.** Body weight changes (ΔW/W₀) before, during, and after partial SD. The gray shaded region indicates the 7-day partial SD. Due to the experimental scheduling, some mice from cohort 2 underwent 9 days of partial SD rather than the standard 7-day protocol, and therefore were excluded from this analysis.
